# Inferring global exponents in subsampled neural systems

**DOI:** 10.1101/2024.11.29.626005

**Authors:** Davide Conte, Antonio de Candia

## Abstract

In systems displaying an activity charaterized by avalanches, critical exponents may give informations on the mechanisms underlying the observed behaviour or on the topology of the connections. However, when only a small fraction of the units composing the system are observed and sampled, the measured exponents may differ significantly from the true ones. In this study, using Branching Process and (2+1)D Directed Percolation we show that some of the exponents, namely the ones governing the power spectrum and the detrended fluctuation analysis (DFA) of the system activity, are more robust and are unaffected in some intervals of frequencies by the subsampling. This robustness derives from the preservation of long-time correlations in the subsampled signal, even though large avalanches can be fragmented into smaller ones. These results don’t depend on the specific model and may be used therefore to extract in a simple and unbiased way some of the exponents of the unobserved full system.

## INTRODUCTION

The evolution of systems observed in Nature is constantly influenced by their response to external perturbations. Numerous studies have demonstrated that many physical systems, despite their macroscopic differences, share similar underlying dynamics. In particular, due to these perturbations, a single event can trigger a cascade of subsequent events, known as an *avalanche*, leading the system to self-organize towards a critical state [1]. This behavior, referred to as crackling noise [2, 3], has been observed across a wide range of phenomena, including earthquakes [4, 5], solar flares [6], biological systems [7, 8], stock markets [9–11], epidemic spread [12] and neural networks [13–15].

An avalanche is typically defined as a chain of consecutive activity occurring above a threshold over an interval of time. Defining *V*(*t*) as the time dependent observable, for example the firing rate in neural network, the interval of time of the avalanche is a consecutive set of time steps such that *V*(*t*) *>* Θ, preceded and followed by two time steps with *V*(*t*) ≤ Θ. The duration *T* of an avalanche is defined as the length of the interval of time characterizing it, while the size *S* of the avalanche is the integral of *V*(*t*) − Θ over the same interval. In the following, we will take the threshold Θ = 0. In crackling noise phenomena, sizes and durations of avalanches follow power-law distributions, which is a defining characteristic of this behavior. The probability distribution of the sizes of the avalanches is given by 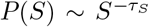, that of durations by 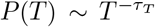, while the average size of avalanches of given duration is ⟨*S*⟩(*T*) ~ *T*^*γ*^. In most cases, these exponents satisfy the relation [2, 16]

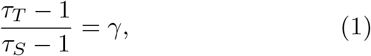

which is generally valid when the system is critical. Relation (1) indeed can be violated when the system is not tuned at criticality, see for example [17, 18].

For a stationary process, in which average quantities do not vary over time, we can define the correlation function of *V*(*t*) as

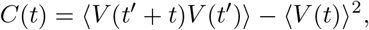

and the power spectrum

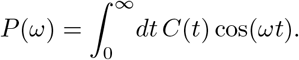

As shown by Kuntz and Sethna [3], this too follow a power law behaviour in crackling noise, e.g. the power spectrum goes as *P*(*ω*) ~ *ω*^−*β*^ at low frequencies, while the high-frequency part is typically dominated by white noise. Note however that, for a finite size system, there is a cut-off in the duration of the avalanches, so that the signal *V*(*t*) becomes uncorrelated at very long times, and another white noise regime is observed at very low frequencies, so that the exponent *β* is actually observed in a finite range of frequencies.

As shown in quite general terms in [3], if different avalanches are uncorrelated, then

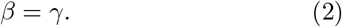

The condition of uncorrelated avalanches is usually well verified when the time separation between avalanches is large, or if the system reaches an absorbing state between different avalanches, so that memory of past activity is lost. The behaviour of power spectrum is strictly connected to that of detrended fluctuation analysis (DFA) [19], that considers the standard deviation of the fluctuations of the detrended signal, over varying intervals of time. For a process *V*(*t*) in discrete time *t* ∈ ℕ, this is defined in the following way: consider an interval of time *t, t* + 1, …, *t* + *n*− 1 of length Δ*t* = *n*, and compute the linear function *Y*_*k*_ = *a* + *bk* that best fits the given function *V*(*t* + *k*) with *k* = 0, …, *n* − 1. Then compute

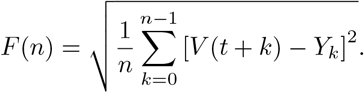

Finally average *F*(*n*) over many non-overlapping intervals of time. The function *F*(*n*) is strictly related to the power spectrum of the function *V*(*t*) [20]. If the power spectrum is proportional to *ω*^−*β*^ in some interval of the frequencies, then *F*(*n*) will be proportional to *n*^*α*^ for a corresponding range of *n*, with *β* = 2*α* − 1.

The previous discussion concerns systems observed in their entirety. However, our knowledge of the system is often limited to a small fraction of it, making it challenging to obtain predictions of the properties of the global system. Different studies [21, 22] have shown that subsampling can significantly alter the observed distributions, even when these distributions are scale invariant. Consequently, the critical exponents measured in sub-sampled data may deviate from those of the fully sampled system, resulting in violations of the scaling relations. In this work, we demonstrate that there exists a frequency (or timescale) range in which the exponents *β* of the power-spectrum exponent, and *α* of the detrended fluctuation analysis, remain invariant with respect to sub-sampling. As the exponent *γ* relating the mean size to the duration instead changes with subsampling, the relation (2) does not hold anymore in the subsampled system. This can be traced back to the fact that, in a subsampled system, subsequent avalanches are not uncorrelated, as they often are successive fragments of a larger avalanche spanning also not sampled regions of the system. While the exponents of avalanche distributions in subsampled systems, e.g. *τ*_*T*_ or *γ*, are computed taking in account those fragments as separated independent avalanches, when computing the power spectrum or the DFA, one considers the temporal correlations of the signal, regardless of the fact that the signal is broken up in successive segments. Such correlations at long time scales are the same as in the fully sampled signal. We demonstrate this property within two particular models, the branching process (mean field directed percolation) and (2+1)D directed percolation. In the first model the property can also be derived analitically, while in the second it is confirmed by numerical simulations. However, from the discussion above it is clear that these results are general and do not depend on the model that produced the signal analysed, being a consequence of the fact that long time correlations are preserved also in the subsampled signal.

This issue has profound implications across various fields. In neural systems, for instance, it is known that the brain operates near a critical state, as this optimizes dynamic range [23], information transmission, and information capacity [24]. Neural activity is therefore characterized by avalanches as in crackling noise, whose critical exponents carry information on the underlying neural dynamics or topology of the network. Current technologies can record neural avalanches only from a small portion of the cerebral cortex, leading potentially to systematic inconsistencies in the values of the measured exponents and in the scaling relations. Developing robust methods to infer the global critical exponents from such partial data is therefore essential. In this context, accurate prediction of critical exponents is particularly valuable, not only for advancing our theoretical understanding of brain function, but also for potentially serving as a diagnostic tool.

## I. BRANCHING PROCESS

The first model we consider is the paradigmatic model of avalanche spreading, that is the branching process. It corresponds to the mean field version of the directed percolation model. We consider a set of *N* nodes of the network, that can be in two states, active and quiescent. Starting from the absorbing state, in which all the nodes are quiescent, we set one node to the active state, to start a new avalanche. Then, at each time step *t* and for each node that is active at that time, we find the nodes that are activated by it at time *t* + 1. We use a all-to-all connectivity, so that each of the *N* nodes can be activated with probability *σ*/*N*, where *σ* is the branching parameter. Nodes that are active at time *t* become quiescent at time *t* +1, unless some node has activated them again. If at some time the absorbing state is reached, the avalanche ends, and at the following time another one is started. In the following we consider the critical value *σ* = 1 of the branching parameter.

To simulate the subsampling of the system, we consider a subset formed by *N*_sub_ < *N* nodes, and analyse the avalanches observed taking in account only the *N*_sub_ nodes. We define the subsampling ratio as *f* = *N*/*N*_sub_, where *f* = 1 means that we observe all the nodes of the system, while larger values correspond to lower density of observed nodes. Due to the all-to-all connectivity, the results are independent of the chosen subset of the nodes, but depend only on the number *N*_sub_ of nodes forming the subset. We measure the distributions *P*_sub_(*S*), *P*_sub_(*T*), and the mean size ⟨*S*⟩ _sub_(*T*) for different subsampling sizes *N*_sub_. We made 256 independent runs, each simulating a system with *N* = 2^24^ ≃1.6 · 10^7^ nodes, and the subsamplig ratio *f* was taken between 1 and 1024. Each run produced about 6 · 10^7^ avalanches for a total time of about 2·10^9^ In Fig. 1 we show the results regarding the distributions of sizes and durations of the avalanches. In panels A, C and E we show respectively *P*_sub_(*S*), *P*_sub_(*T*) and ⟨*S*⟩ _sub_(*T*) for the different subsampling ratios, while in panels B, D and F we show the corresponding logarithmic derivatives. Given a function *f*(*x*) ∝ *x*^*α*^, the logarithmic derivative is defined as 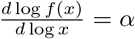, so that it gives a measure of the effective exponent governing the function around some value of the variable. A range of values where the logarithmic derivative is nearly constant corresponds to a range where the function can be well approximated by a power law. As can be seen in Fig. 1B, the exponent of the distribution of the sizes in the fully sampled system is equal to 3/2 in the whole interval of sizes, except near the cut-off of the distribution. For sub-sampling ratios between 4 and 256, one can see larger and larger deviations for small sizes, that correspond to the first steeper decay of the distribution *P*(*S*). (These are called the “hairs” of the distribution in Ref. [21]). There remains however a smaller and smaller interval of sizes that display the same exponent 3/2 of the full system. Only for subsampling ratio 1024 the distribution always displays a larger exponent (a steeper decay).

**FIG. 1.**
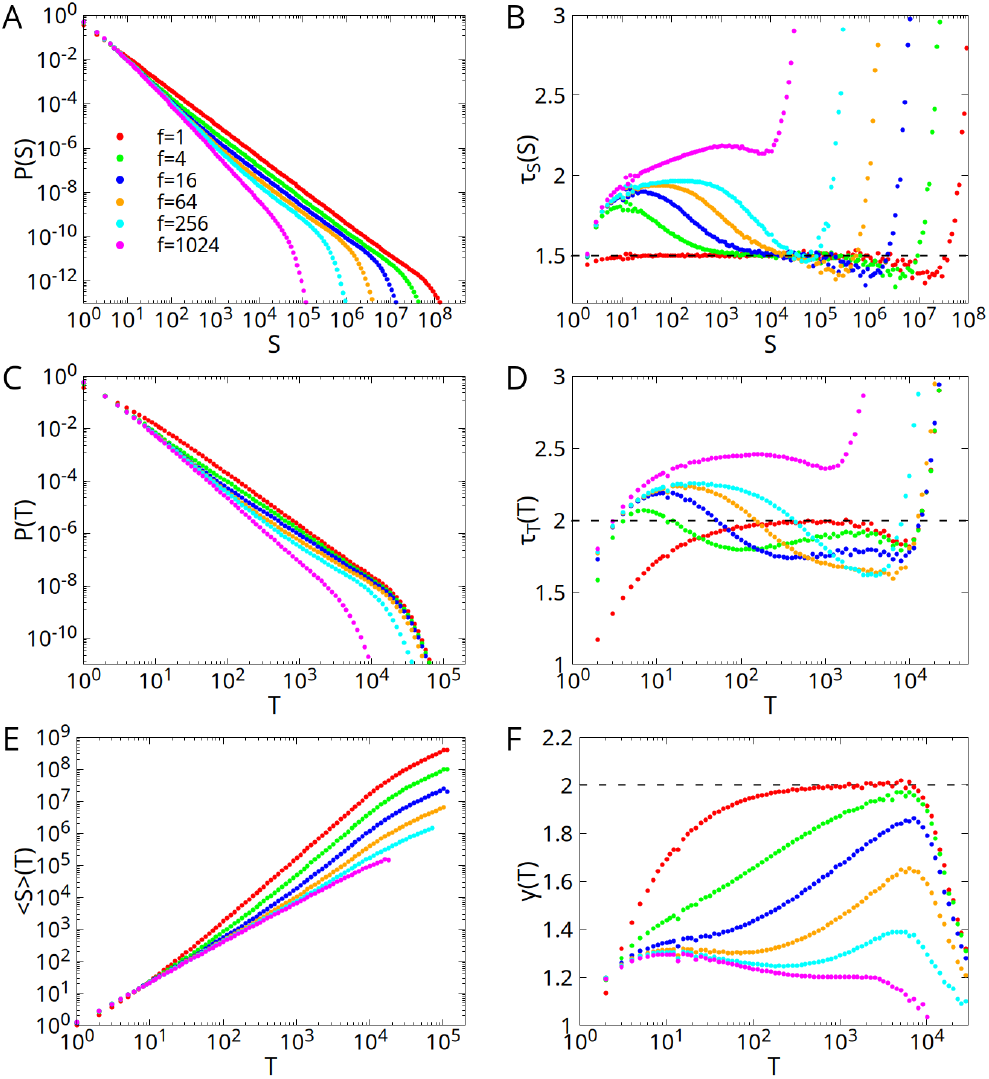
Avalanches distributions for the branching process on a lattice with *N* = 2^24^ sites. Legend shows different sub-sampling ratios *f* = *N/N*_sub_. Note that *f* = 1 corresponds to the full lattice. Left panels A, C, and E: distributions of the sizes, durations, and mean size as a function of the duration. Right panels B, D, and F: logarithmic derivatives of the functions on the left. A range of values where the logarithmic derivative is nearly constant correspond to a range where the function on the left can be approximated by a power law. Dashed lines represent the expected exponents for the fully sampled branching process.

The distributions of the durations on the other hand have a different behaviour. A large deviation from the expected exponent 2 is observed for small durations already in the fully sampled case (red dots in Fig. 1D). For sub-sampled systems, the exponent 2 is never observed for any range of the duration. Instead, the functions have an initial steeper decay (a larger exponent), and then for large durations a range with an exponent smaller than 2. Also the value of the exponent *γ*, that is related to both the exponents *τ*_*S*_ and *τ*_*T*_, is different from the expected value 2 for all the intervals of duration, and hovers around the value *γ* ≃1.3 for example in the case of a subsampling 256. Therefore, we can conclude that, while there is a range of sizes where the exponent *τ*_*S*_ remains stable also for subsampled systems, exponents *τ*_*T*_ and *γ* have strong deviations that make difficult to infer the values for the global system.

We now show that the exponent governing the power spectrum remains unchanged for a range of frequencies in the case of the subsampled systems. To do this, let *N* and *V*(*t*) be respectively the total number of sites and the number of active sites at time *t* in the complete lattice, *N*_sub_ and *V*_sub_(*t*) the corresponsing quantities in the “sampled” set of sites, that is a subset (*N*_sub_ < *N* and *V*_sub_(*t*) ≤ *V*(*t*)) of the complete lattice. Due to the all-to-all connectivity, at each step only the number *V*(*t*) of the active sites depend on the state of the system at time *t* − 1, but not the particular subset of sites that is activated. Therefore, the probability that *V*_sub_(*t*) of the sites of the sampled sublattice are activated is the probability of extracting *V*_sub_(*t*) active sites, in *N*_sub_ random draws without replacement, from a population of *N* sites that contains exactly *V*(*t*) active sites, that is the hypergeometric distribution [25]

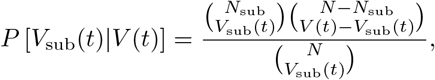

If *P* [*V*(*t*)] is the probability of having *V*(*t*) active sites in the complete lattice at time *t*, and *P* [*V*(*t*), *V*(*t*′)] the probability of having *V*(*t*) and *V*(*t*′) active sites at times *t* and *t*′, then the expected values of *V*_sub_(*t*), *V*_sub_(*t*)^2^ and *V*_sub_(*t*)*V*_sub_(*t*′) are

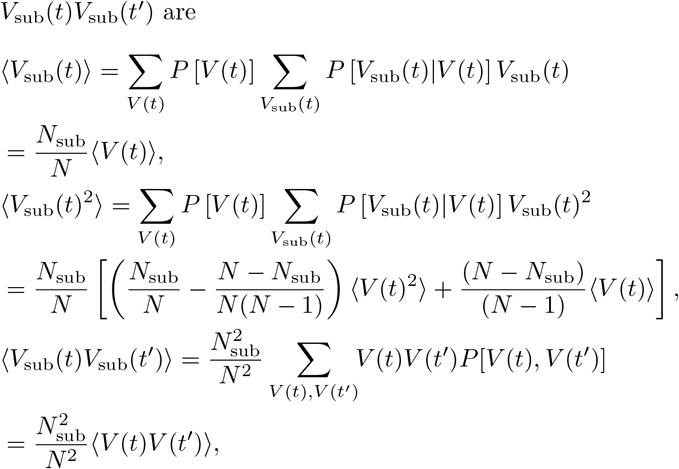

where the last equation holds for *t* ≠ *t*′. Therefore, the correlation function of *V*_sub_(*t*) is given by

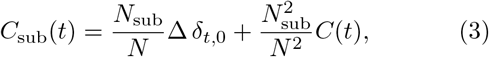

where *C*(*t*) is the correlation function of *V*(*t*), and

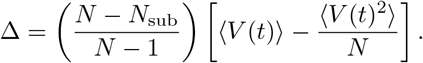

Note that for *N*_sub_ ≪ *N* the coefficient Δ does not depend on the subsampling ratio *N*/*N*_sub_. Defining the power spectrum *P*_sub_(*ω*) as the Fourier transform of the subsampled correlation function *C*_sub_(*t*), we have

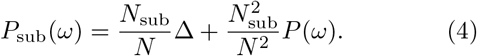

The first term corresponds to a white noise, due to the very fast decorrelation induced by the random choice of the *V*_sub_(*t*) active sites at each step, which is a characteristic of the branching process on a fully connected lattice. Apart from that term, the power spectrum of the number of active sites in the subsampled lattice is exactly proportional to the power spectrum of the full lattice. At the critical point, where *P*(*ω*) = *kω*^−*β*^ with *k* a constant and *β* = 2, the subsampled power spectrum *P*_sub_(*ω*) will be proportional to *ω*^−*β*^ for frequencies *ω* < *ω*^*^, and constant for higher frequencies, where 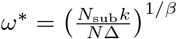.

In Fig. 2 we show the results of the simulations of the branching process on the fully connected lattice with *N* = 2^24^ sites. In panel A we show the power spectrum *P*_sub_(*ω*) of the number of active sites on the sampled sublattice. The results confirm the expression found before, Eq. (4). Note that at very low frequencies there is a white noise plateau, due to the finite size of the lattice and therefore to the cut-off in the distribution of avalanche durations. For times longer than the cut-off (and frequencies lower than the inverse cut-off) the signal we are analysing is decorrelated because there are no avalanches during such a long time.

**FIG. 2.**
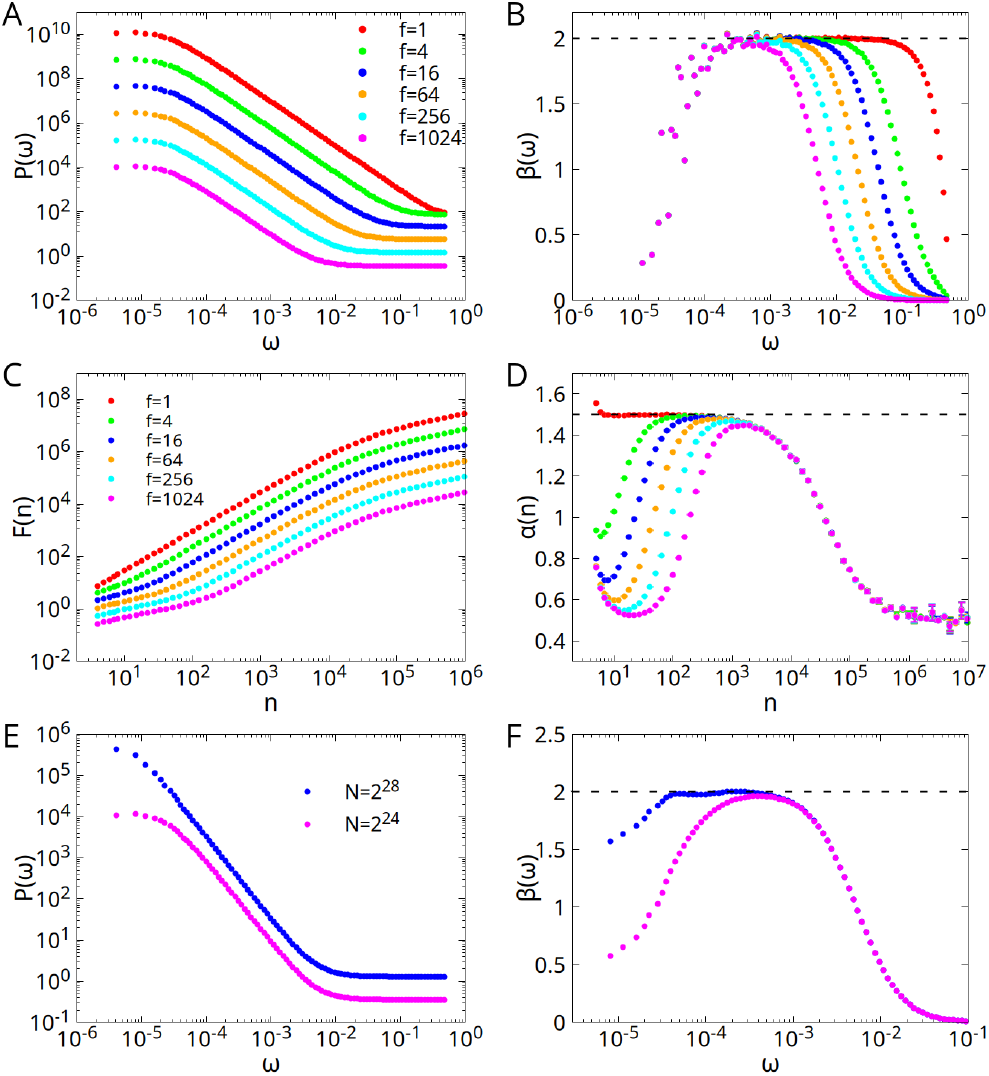
Panels A and C: power spectrum and detrended fluctuation analysis of the signal *V*(*t*) or *V*_sub_(*t*), the number of active sites belonging to the whole lattice or to the sampled subset of the lattice, for various subsampling ratios. Panels B and D: logarithmic derivatives of the functions on the left. Panels E and F: power spectrum and its logarithmic derivative for subsampling ratio *f* = 1024 and for two different sizes. Data in panel F have been filtered to reduce statistical fluctuations.

We also consider the detrended fluctuation analysis (DFA) of the signal [19], previously defined. In Fig. 2C we show the measured functions *F*(*n*) and *F*_sub_(*n*) for *V*(*t*) and *V*_sub_(*t*). As in the case of the power spectrum, the data always display an interval of frequencies, or interval lenghts, where the exponent in the case of subsampled data is equal to that of the fully sampled lattice, see Fig. 2D. This invariance can be therefore exploited to infer exponents of the whole system using only subsampled data. Note that, the larger the subsampling ratio, the smaller the frequency where the high frequency white noise regime sets in. Therefore, for large subsampling ratios, the range where the plateau in the value of the exponent is found may be small, being limited by the two white noise regimes at high and low frequency. However, for larger system sizes, the low frequency regime shifts towards lower frequencies, therefore enlarging the intermediate plateau where the correct exponent is found. This is shown in panels E and F, where the power spectrum and its logarithmic derivatives are shown for the same subsampling ratio *f* = 1024 and for two sizes *N* = 2^24^ and *N* = 2^28^. In the larger system the range where the expected exponent *β* = 2 is found is correspondingly larger.

## II. DIRECTED PERCOLATION ON A (2+1)D LATTICE

In this Section, we consider the directed percolation model on a (2+1)D lattice, that corresponds to a branching process not on a fully connected lattice but on a 2D lattice. While the critical exponents in this case are different from the mean field values found in the previous Section, we again find that power spectrum and detrended fluctuation exponents remain invariant in the case of subsampling for a range of frequencies or interval lengths.

The directed percolation model on a (2+1)D lattice is defined as follows. Consider a 2D square lattice, where each site can be activated or quiescent. Each site that is activated at time *t* may activate at time *t* + 1 each of its four neighbors with probability *p*. A node that is active at time *t* becomes quiescent at time *t* + 1, unless some of its neighbors has activated it again. As in the case of the branching process studied in the previous Section, if all the sites are quiescent, we start a new avalanche selecting a site at random and activating it. We simulate the system at the critical value of the probability, *p*_*c*_ ≃0.28729 [26], and for *L* = 1024, *N* = *L*^2^ ≃ 10^6^ sites, until 10^7^ avalanches have been observed.

To assess the effects of subsampling also in the case of (2+1)D directed percolation, we take a subset of the sites of the lattice as done in the previous case, and consider the number *V*_sub_(*t*) ≤*V*(*t*) of sites active at time *t*. In Fig. 3 we show the criterion used to choose the subset of sites for subsampling ratios 2, 4 and 8. In a similar way sites are selected for higher ratios. We have considered ratios between 1 and 1024.

**FIG. 3.**
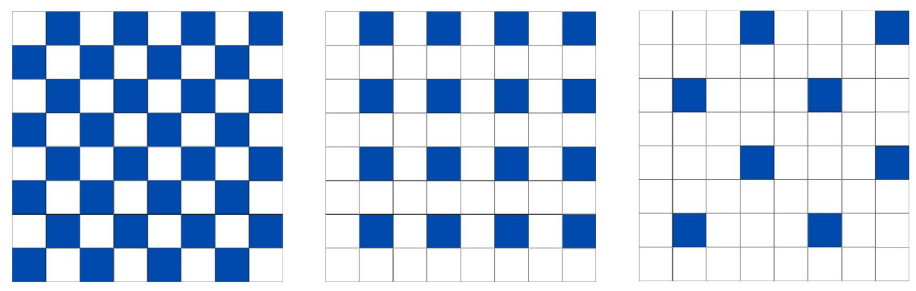
Blue sites are the ones selected for subsampling, for ratios *N/N*_sub_ = 2, 4, 8. In a similar way sites are selected for higher ratios.

To check also the dependence on temporal binning, we also use different temporal bin sizes. Specifically, for a bin size *w* = 2, 4, …, we define a new function *V*_sub,*w*_(*t*), with *t* ∈ ℕ, as the sum of *V*_sub_(*t*′) for *t*′ = *wt, wt*+1 … *w*(*t*+1) − 1, and then compute avalanche distributions and power spectrum for this new function. Note that, in the subsampled case, it is essential to choose a temporal bin *w* = 2 or larger. This is because, for the subsampling criterion chosen, sampled (blue) sites can be active only every other time step, so that with *w* = 1 every avalanche would be divided into pieces of duration one step. We considered temporal bins between *w* = 2 and *w* = 64. Given the number *V*(*t*) or *V*_sub_(*t*) of active sites as a function of time, consecutive intervals with *V*(*t*) > 0 or *V*_sub_(*t*) > 0 correspond to avalanches. In Fig. 4 we show the distributions of sizes, durations, mean size versus duration, and the power spectrum for different subsampling ratios. Each of the distributions exhibits a power-law behavior in a large range, and we extract exponents from a least squares fit in that range, excluding the cut-off due to the finite size of the lattice. In the case of the fully sampled lattice, we obtained the following exponents: *τ*_*S*_ = 1.27 ± 0.03, *τ*_*T*_ = 1.46 ± 0.01, *γ* = 1.69 ± 0.02, *β* = 1.70 ± 0.01, consistent with those found in the literature [27], and satisfying the scaling relations of crackling noise (1) and (2).

**FIG. 4.**
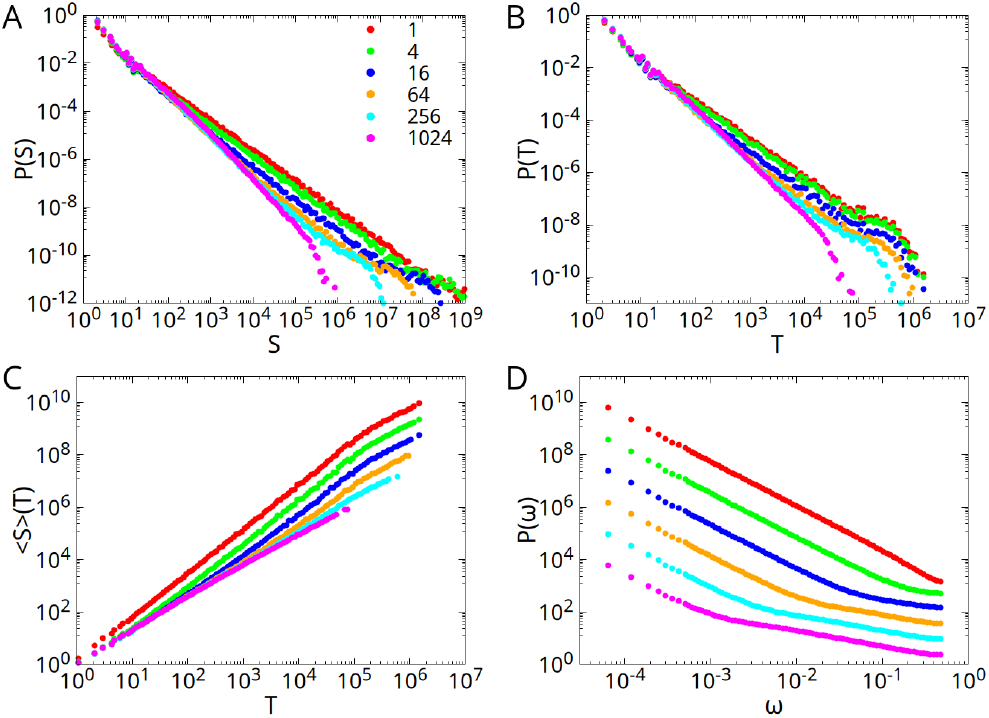
Avalanche distributions and power spectrum for different subsampling ratios in (2+1)D directed percolation, for a temporal bin *w* = 2. We estimated the critical exponents, along with their corresponding errors, using a linear fit applied only to the interval with a linear trend.

As in the previous case, by applying subsampling, we expect variations in the probability distributions and, consequently, in the respective critical exponents. The effect of subsampling is to fragment a single avalanche into smaller avalanches with reduced size and duration, thus leading to an increase of the exponents for the probability distributions of sizes and durations. This is shown in Fig. 6A and B, that reports the exponents of the avalanche distributions as a function of the sampling size, for different temporal binnings. As shown in Fig. 6C, the resulting effect of subsampling on the mean size ⟨*S*⟩ _sub_(*T*) as a function of duration, is to progressively reduce the exponent as the sampling size decreases, as already observed in the case of the mean field branching process.

As already observed in [28], the effects of subsampling can be reduced by increasing the width of temporal bins, so that different fragments of the same avalanche are reconnected again. Therefore, the values of the exponents for the fully sampled system tend to be recovered. This is confirmed by our data, where exponents for larger temporal binning remain stable up to higher values of the subsampling ratios.

In Fig. 6D, we show the exponent *β* of power spectrum decay as a function of subsampling, for different temporal binning. While we have seen that the exponent *γ* varies under sampling procedures, as shown in Fig. 6C, the exponent *β* of the power spectrum remains unchanged, as in the case of the branching process. This is because the power spectrum is the Fourier transform of the temporal autocorrelation function between sites within the same avalanche. Although sampling fragments the avalanche into smaller ones, these fragments are not independent but correlated, as they still belong to the same avalanche propagating through the lattice. On one hand, this leads to the violation of the relationship (2), that holds only in the case of uncorrelated avalanches [3]. On the other hand, the exponent *β* remains unchanged under subsubsampling, while *γ* varies, as shown in the inset of Fig. 6D.

The relation between the correlation function of *V*_sub_(*t*) and that of *V* (*t*) cannot be evaluated explicitly as in the case of the mean field branching process. However, the form of the power spectrum shows that the correlation function, which is its Fourier antitransform, with respect to Eq. (3) generalizes to

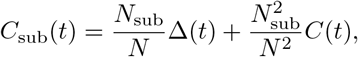

where Δ(*t*) does not depend on 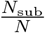 if *N*_sub_ ≪ *N*, and decays in a short time. It represents the excess probability (with respect to a random choice of sites) that the avalanche at time *t* overlaps with the same sites of the sampled set as those at time 0. Therefore, the power spectrum in the subsampled case will exhibit two different behaviours: at low frequencies it will be given by 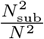 times the power spectrum of the fully sampled case. At high frequencies on the other hand, it will go as the Fourier transform of Δ(*t*). The plots of Fig. 4D confirm this prediction. While at low frequencies the spectra all follow a power law with exponent *β* ≃ 1.7, at high frequency on the other hand they follow a different power law, with an exponent β_2_ ≃ 0.6.

Consistent with previous findings for the branching process, the detrended fluctuation analysis (DFA) also exhibits power-law scaling across varying sampling ratios, as illustrated in Fig. 5A. Notably, the observation of a distinct critical exponent *β*_2_ for the power spectrum is similarly observed in the DFA: as sampling density increases, a new linear trend emerges for small values of *n*, with an exponent around 0.8. This result aligns with the power spectrum, as the two exponents are connected through the scaling relation *β* = 2*α* − 1. Fig. 5B further demonstrates that the exponent *α* remains approximately invariant across different sampling ratios and temporal binning, consistently satisfying the previously stated scaling relation and the results observed for the branching model.

**FIG. 5.**
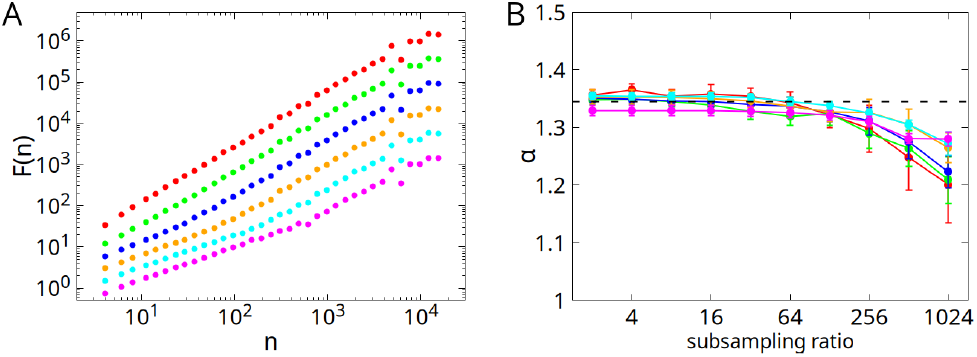
Left: detrended fluctuation analysis of the subsampled active sites for different subsampling ratios and bin size *w* = 2. Right: critical exponent *α*, remains approximately invariant during subsampling procedures, confirming, within error margins, the relationship with the *β* exponent of the power spectrum *β* = 2*α* − 1.

**FIG. 6.**
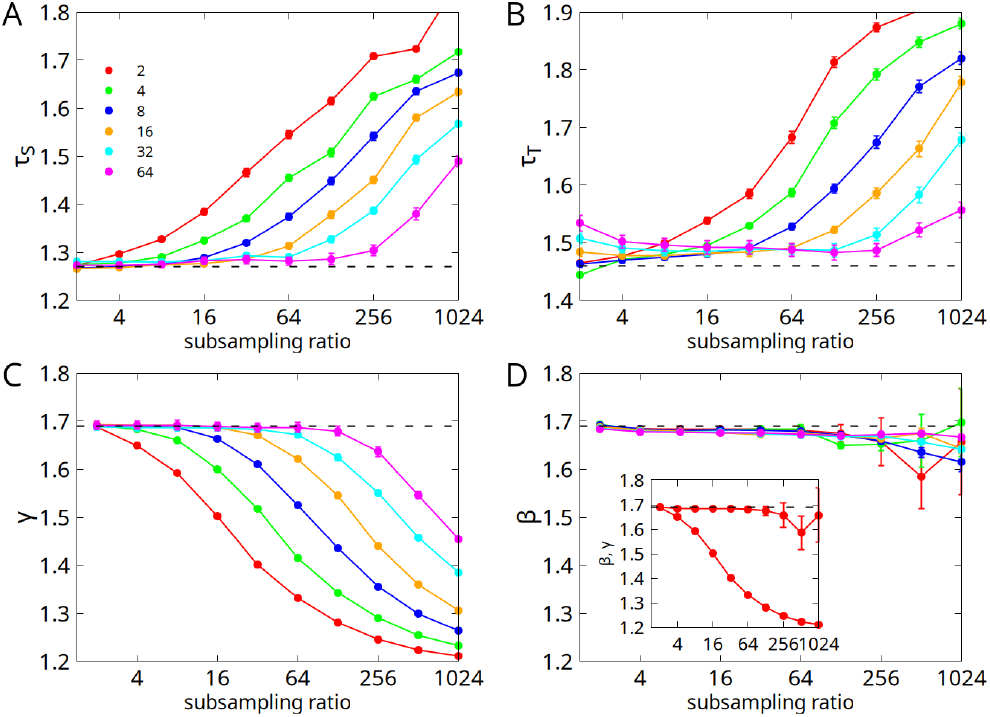
In the figure the critical exponents *τ*_*S*_, *τ*_*T*_, *γ* and *β*, extracted with a least squares fit as in Fig. 4, are shown for different subsampling ratios and different temporal binning. We observe that for the exponents *τ*_*S*_ and *τ*_*T*_ of *P*_sub_(*S*) and *P*_sub_(*T*), the effect of subsampling results in an increase in the value of the critical exponent, due to the fragmentation of large avalanches into smaller ones. The exponent *γ* of ⟨ *S*⟩ _sub_(*T*) instead decreases with the subsampling. As observed in [28], increasing the temporal binning allows the global value of the exponent to be recovered. On the other hand, the exponent *β* of the power spectrum remains nearly unchanged under subsampling, and for any value of the temporal binning. Inset of panel D: Comparison between the exponent of the power spectrum *β*, and that of the average avalanche size *γ*, as a function of the subsampling ratio, for a bin size *w* = 2. The two exponents coincide within errors for the case of the complete lattice, both close to 1.7, but as the number of sampled sites decreases, the discrepancy between them increases. Specifically, *β* remains nearly unchanged across subsampling procedures, while *γ* decreases.

## III. CONCLUSIONS

Inferring global critical exponents in systems like neural networks has been a longstanding challenge. Since the seminal work of Beggs and Plenz [13], it has been hypothesized that the critical exponents of neural avalanches would align with those of the branching process (BP). However, recent advances in technology and research have revealed the existence of critical exponents that differ from those of the BP universality class, suggesting the possibility of multiple universality classes in neural networks [29–32].

Given that studies on neural avalanches are typically based on sampled portions of the system, accounting for subsampling effects is crucial. Previous work by Levina and Priesemann [21] has shown that subsampling can distort avalanche distributions, potentially leading to inaccurate predictions of global critical exponents.

In this study, using two stochastic models, the branching process and (2+1)D directed percolation, we have demonstrated that the exponentes *β* of power spectrum and *α* of detrended fluctuation analysis (DFA) are robust with respect to subsampling. Given that these exponents are related to the exponent *γ* of size versus duration of the avalanche in the fully sampled case, they can be used to predict the latter when only subsampled data are available.

The underlying reason for this robustness is that the long-time correlations in the activity persist under sub-sampling: although a single large avalanche may be fragmented into multiple smaller ones, these pieces still form a correlated chain in time. Consequently, the low-frequency region of the power spectrum and the long-range correlations detected by DFA remain approximately unchanged. In contrast, exponents of avalanche-size or avalanche-duration distributions, such as *τ*_*T*_ or *γ*, treat those fragments as independent events and are therefore strongly affected by subsampling, leading to the violation of the crackling noise relation *β* = *γ*, which pre-supposes uncorrelated avalanches.

Although we have demonstrated these findings for two specific models, the preservation of long-range temporal correlations in partially observed avalanche processes suggests that this effect is not confined to those models alone. Critical exponents may indeed differ for other network topologies, or under different dynamical processes, but the arguments about fragmentation of avalanches versus preserved time correlations apply quite broadly. In the case of the brain, accurate predictions of these exponents could be crucial for identifying variations in the neural activity or in the network topology, and therefore neurological disorders or pathologies. Moving forward, our goal is to extend the findings from this study, demonstrated for the branching process and directed percolation, to more realistic models, ultimately contributing to a deeper understanding of criticality in neural systems.

